# TUDCA treatment restores aortic and perivascular adipose tissue function in post-weaning protein-restricted mice

**DOI:** 10.64898/2026.03.11.711218

**Authors:** Israelle N Freitas, Carolina M Lazaro, Joel A da Silva Junior, Kenia M de Oliveira, Jamaira A Victorio, Everardo M Carneiro, Ana P Davel

## Abstract

**Background:** Early-life protein restriction is a risk factor for cardiovascular disease, yet the mechanisms underlying vascular dysfunction and therapeutic strategies remain poorly defined. Tauroursodeoxycholic acid (TUDCA) is a bile acid that inhibits endoplasmic reticulum (ER) stress and has therapeutic potential in metabolic diseases. We hypothesized that TUDCA exerts vasculoprotective effects in the setting of post-weaning protein restriction.

**Methods:** Post-weaning male and female mice fed a normoprotein (14% protein) or protein-restricted (6% protein, isocaloric) diet for 105 days. In the last 15 days, mice received TUDCA (300 mg/kg/day) or vehicle. Vascular function was assessed in the thoracic aorta with or without perivascular adipose tissue (PVAT). mRNA expression and histological analyses were performed in aorta and PVAT.

**Results:** Long-term protein restriction resulted in endothelial dysfunction, vascular hypocontractility, and loss of the anticontractile effect of PVAT in males, but not females. These alterations were restored by TUDCA. In aorta, TUDCA normalized expression of eNOS and contractile phenotype-related genes α-actin, SM22α, Cav1.2 whereas, in the PVAT, TUDCA restored lipid content and expression of PRDM16, PPARγ, PGC1α, leptin, and OB-Rb in protein-restricted mice. TUDCA attenuated fibrosis and ER stress markers while increased the bile acid receptor FXR expression in both tissues. Similar to TUDCA, ER stress inhibition with 4-phenylbutyric acid restored vascular and PVAT function in protein-restricted male mice.

**Conclusions:** Post-weaning protein restriction induces vascular and PVAT dysfunction and fibrosis in males, associated with ER stress. TUDCA significantly attenuates these alterations, supporting its potential as a therapeutic strategy for vascular complications associated with early-life undernutrition.

## INTRODUCTION

Malnutrition refers to deficiencies or excesses in nutrient intake, imbalances of essential nutrients, or impaired nutrient utilization. In 2024, the estimated prevalence of undernourishment was 8.2%, a number that has risen notably since 2019, further sharpened by the COVID-19 pandemic (1). In addition, there are marked regional differences in the prevalence of undernourishment, such as subregions of Africa and South Asia had approximately one third of all children affected by stunting (2). Even higher rates have been reported among the Yanomami Indigenous population in Brazil, where in 2022, 52.2% of children under 5 years of age were undernourished (3). Early-life undernutrition may have long-term health consequences, including an increased risk of cardiovascular disease in adulthood. However, therapeutic strategies to mitigate the long-term consequences of undernutrition remain poorly defined.

Early-life protein restriction contribute to metabolic and cardiovascular abnormalities increasing the susceptibility to chronic diseases (4). In agreement, increased blood pressure and endothelial dysfunction was demonstrated in rodent models of low protein diet in early stages of development (5–7). In early stages of development, protein restriction might increase perirenal, perigenital and visceral fat thereby programming susceptibility to obesity with increased expression genes involved in lipogenesis and adipogenesis (8; 9). But no changes in adiposity were observed in response to a postnatal low-protein diet (10; 11). Therefore, dietary protein restriction may affect adiposity depending on the stage of development and the fat depot. The perivascular adipose tissue (PVAT) is an ectopic fat depot surrounding most blood vessels regulating vascular tone and structure, and blood pressure and a dysfunctional PVAT contribute to vascular disorders (12; 13). Although PVAT has been considered an important player in cardiovascular homeostasis, the impact of different forms of undernutrition on PVAT function and structure remains unknown.

Endoplasmic reticulum (ER) stress signaling is a pathophysiological mechanism involved in several metabolic and cardiovascular diseases including obesity, type 2 diabetes, non-alcoholic steatohepatitis (NASH), and hypertension (14; 15). ER stress has been associated with dysfunctional white and brown adipose tissues during obesity that exhibit enhanced expression of unfolded protein response (UPR) markers such as activated transcription factor-6 (ATF-6), activated transcription factor-4 (ATF-4), and C/EBP Homologous Protein (CHOP) (16; 17). In PVAT, ER stress has been associated with vascular dysfunction, vascular inflammation, and atherosclerotic plaque destabilization (18–20). In the setting of dietary protein deficiency, a decrease in protein turnover could be related to ER stress (21) but remains to be determined if this mechanism is involved in vascular and PVAT alteration.

The bile acid tauroursodeoxycholic (TUDCA) is a US FDA-approved hydrophilic bile acid for the treatment of chronic cholestatic liver disease. Beyond its classical digestive function, it has chemical chaperone activity thereby inhibiting ER stress. Therefore, TUDCA has emerged as a therapeutic strategy to minimize metabolic alterations associated with obesity and type 2 diabetes (18; 22; 23). This agrees with reduced adiposity, improved glucose homeostasis and insulin signaling in adipose tissue of protein-restricted mice exposed to high-fat diet (11). TUDCA effects may also be associated to cell signaling through receptors including the farnesoid X receptor (FXR), the G protein-coupled bile acid receptor (TGR5) and sphingosine-1-phosphate receptor 2 (S1PR2). These receptors are classically expressed in the liver and digestive tract but have been also identified in blood vessels and adipocytes (24; 25). In this study, we hypothesized that TUDCA treatment would improve vascular and PVAT function and structure in the setting of early-life protein restriction. We found that TUDCA mitigates protein restriction-induced endothelial and PVAT dysfunction positioning ER stress inhibition as a target mechanism to mitigate vascular complications associated with protein restriction.

## METHODS

### Data Availability

The data that support the findings of this study are available from the corresponding author on reasonable request. A detailed description of all methods is in the Supplemental Material.

### Animals

All procedures were in accordance with the Ethical Principles of the National Council for the Control of Animal Experimentation (CONCEA/Brazil). Female and male C57BL6/JUnib mice, aged 28 days (post-weaning) received either a normoprotein diet (N; 14% protein, 3.81 kcal/g) or a restricted protein diet (R; 6% protein, 3.81 kcal/g) for 105 days. In the last 15 days, mice were randomly assigned to receive daily PBS (vehicle), TUDCA (300 mg/kg, i.p.) (26) or 4-phenylbutyric acid (PBA) (300 mg/kg, i.p.) (27).

### Blood Pressure

Systolic blood pressure (SBP) was measured via plethysmography.

### Vascular Reactivity

The thoracic aorta with or without adjacent perivascular adipose tissue (PVAT+ or PVAT-, respectively) were placed in an organ bath system to evaluate contractile responses to KCl and phenylephrine or endothelium-relaxation responses to acetylcholine. The maximum effect (Emax) and potency (-LogEC_50_) to agonists were calculated (GraphPad Prism 8.0).

### Histological Analysis

Paraffin-sections (5 µm) from transverse segments of the thoracic aorta and its PVAT were obtained and stained with hematoxylin and eosin and Masson’s trichrome. Images were captured using an optical microscope at 20x magnification.

### Gene Expression

qRT PCR was used for mRNA expression analysis from the thoracic aorta and its respective PVAT. Real-time PCR was performed in 7500 Fast Real-Time PCR System (Applied Biosystems, California, USA) using SYBR® Green Taq ReadyMix™ (Sigma-Aldrich, Saint Louis, USA) or Fast SYBR® Green PCR Master Mix (Applied Biosystems, Massachusetts, USA). Primer sequences are shown in Table S1.

### Statistics

Data were analyzed using GraphPad Prism® 8 software and are presented as mean ± standard error (SE). Outliers were identified and removed using the Grubb’s test (alpha = 0.05). The data normality was tested by the Shapiro-Wilk test. Student’s *t*-test was used for comparing two groups, while one and two-way ANOVA with Tukey post-test was employed for three or more groups, or the respective non-parametric test. A significance level of P≤0.05 was considered statistically significant.

## RESULTS

### TUDCA reverses early-life protein restriction-induced higher blood pressure and impaired vascular reactivity in male mice

Post-weaning protein restriction reduced body weight and total plasma protein concentration and increased blood pressure without changes in fasting glycemia in male (Table 1) and in female mice (Table S2). Consistent with previous reports (6; 7), in males, post-weaning protein restriction impaired endothelial function, as shown by reduced acetylcholine-induced relaxation (Figure 1A, Table S3). In addition, contraction responses to phenylephrine and KCl were reduced in aorta from R compared with N group (Figure 1B, Table S3). Impaired aortic reactivity in the R group was associated with reduced gene expression of vascular smooth muscle contractile phenotype markers including α-actin, SM22α, α1 subunit of the L-type calcium channel, and eNOS (Figure 1C). In females, early-life protein restriction did not affect vascular reactivity (Figure S1A-B). Therefore, TUDCA treatment was conducted in males only. TUDCA treatment did not impact body weight, plasma protein concentration or blood glucose, but normalized the blood pressure (Table 1), as well as the aortic contraction and relaxation responses in male R group (Figure 1, Table S3).

**Figure 1.**
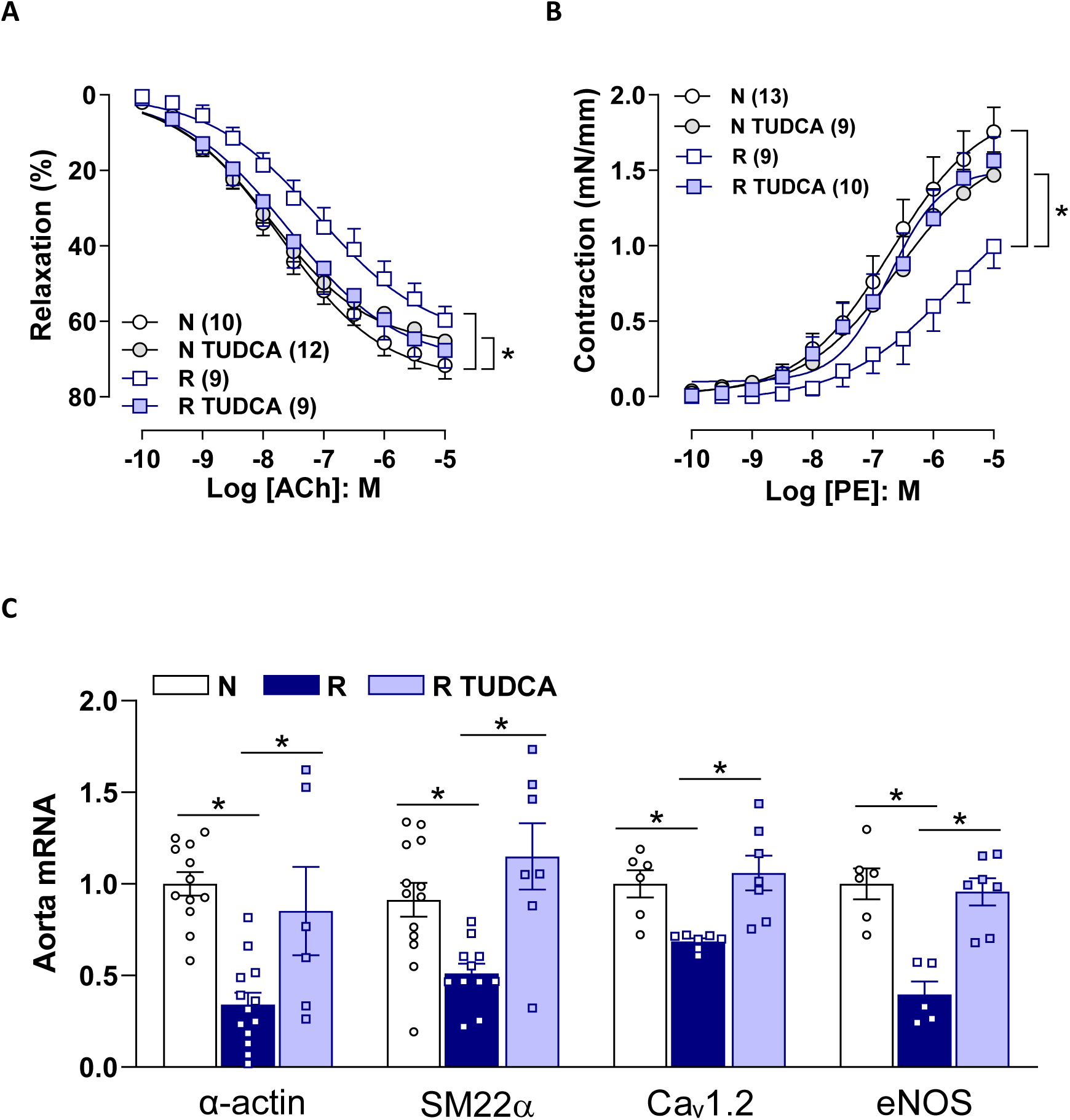
TUDCA treatment ameliorated endothelium-dependent relaxation and vascular contractility in the aorta of protein-restricted mice. Concentration-response curves for acetylcholine (ACh) **(A)** and phenylephrine (PE) **(B)** in aortas from mice fed a normoprotein (N) or protein-restricted (R) diet, with or without TUDCA treatment. Aortic mRNA expression of α-actin, SM22α, Caᵥ1.2, and eNOS **(C)**. Aortic mRNA expression was normalized to Cypa. Data are presented as mean ± SE. *P<0.05: 2-way (A-B) or 1-way ANOVA (C).

**Table 1:** General characteristics of male mice fed a normal-protein (N) or protein-restricted (R) diet, with or without TUDCA treatment.

|  | N<br>(14) | N TUDCA<br>(12) | R<br>(9) | R TUDCA<br>(10) |
| --- | --- | --- | --- | --- |
| Body Weight (g) | 28 ± 2 | 28 ± 1 | 25 ± 5* | 26 ± 1<sup>§</sup> |
| Total Plasma Protein (g/dL) | 6.5 ± 1.0 | 6.6 ± 1.0 | 4.8 ± 0.8* | 5.4 ± 1.4* |
| Blood Pressure (mmHg) | 101 ± 4 | 101 ± 6 | 110 ± 6* | 101 ± 6<sup>#</sup> |
| Blood Glucose (mg/dL) | 99 ± 8 | 97 ± 6 | 99 ± 7 | 97 ± 6 |

Data are mean ± SE; number of animals (n) in each group is in parenthesis.
2-way ANOVA, P≤0.05: \*R vs. N; <sup>§</sup>R TUDCA vs. N TUDCA; <sup>#</sup>R TUDCA vs. R.

### Impaired anticontractile effect and remodeling of PVAT in aorta of protein-restricted mice are restored by TUDCA treatment

PVAT exerts an anticontractile effect in the thoracic aorta of both male and female mice, which may become dysfunctional in obesity in both sexes (revised in 28). Here, we investigated if PVAT anticontractile function is involved in the vascular effects of protein restriction. As expected, phenylephrine-induced contraction was attenuated by PVAT in both male and female aorta of N group (Figure 2A, Figure S1C-D). This effect of PVAT was lost in male R group (Figure 2B, Table S3) but was intact in female R group (Figure S1C-D). This indicates that early-life protein restricted diet affects the anticontractile effect of aortic PVAT in a sex-specific manner. TUDCA treatment restored the anticontractile effect of PVAT in response to phenylephrine in aorta of male R group (Figure 2A-B, Table S3).

**Figure 2.**
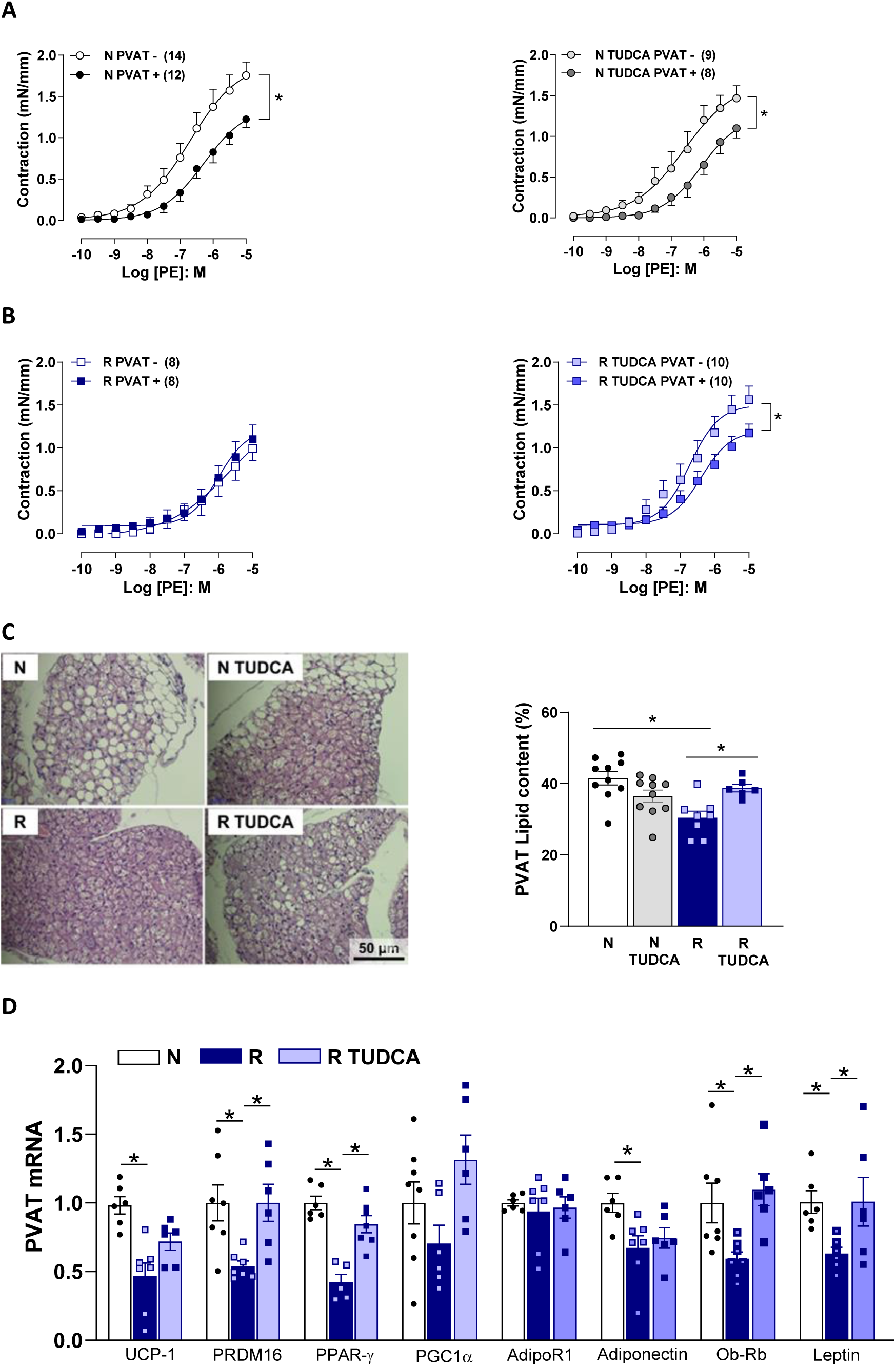
The altered anticontractile effect, lipid content and thermogenic and adipogenic markers expression in the PVAT from protein-restricted mice were restored by TUDCA. Effect of PVAT (+) on the concentration-response curves for phenylephrine (PE) in aortas from mice fed a normoprotein (N) or protein-restricted (R) diet without (**A**) or with TUDCA treatment (**B**). Morphology of aortic PVAT adipocytes assessed by HE staining (bar= 50 µm) for quantification of fat **(C)**. PVAT mRNA expression of UCP-1, PRDM16, PPARg, PGC1α, receptor AdipoR1, adiponectin, receptor Ob-Rb and leptin **(D)**. PVAT RNA expression was normalized to Rplp0 as internal control. Data are presented as mean ± SE. *P<0.05: 2-way (A-B) or 1-way ANOVA (C-D).

PVAT of the thoracic aorta is characterized by a high prevalence of brown adipocytes, expressing thermogenic and adipogenic genes (29). In the R group, we observed reduced PVAT lipid content (Figure 2C) associated with lower gene expression of the thermogenic marker UCP1, the adipogenic markers PRDM16 and PPARy, as well as the vasodilator factors adiponectin and leptin, and its receptor Ob-Rb (Figure 2D). TUDCA restored PVAT lipid content (Figure 2C), accompanied by normalization of PRDM16, PPARγ, Ob-Rb and leptin, without altering UCP-1, PGC1-α, adiponectin and its receptor AdipoR1 in protein-restricted mice (Figure 2D).

Vascular and PVAT fibrosis impacts vascular structure and stiffness (30). Collagen deposition was increased in the aorta and in its PVAT of male R group compared to the N group, which was reversed by TUDCA treatment (Figure 3A-B). This agrees with an increased gene expression of collagen IV in aorta and collagen III and TGFβ in both aorta and PVAT of R group, that were normalized by TUDCA (Figure 3C-D).

**Figure 3.**
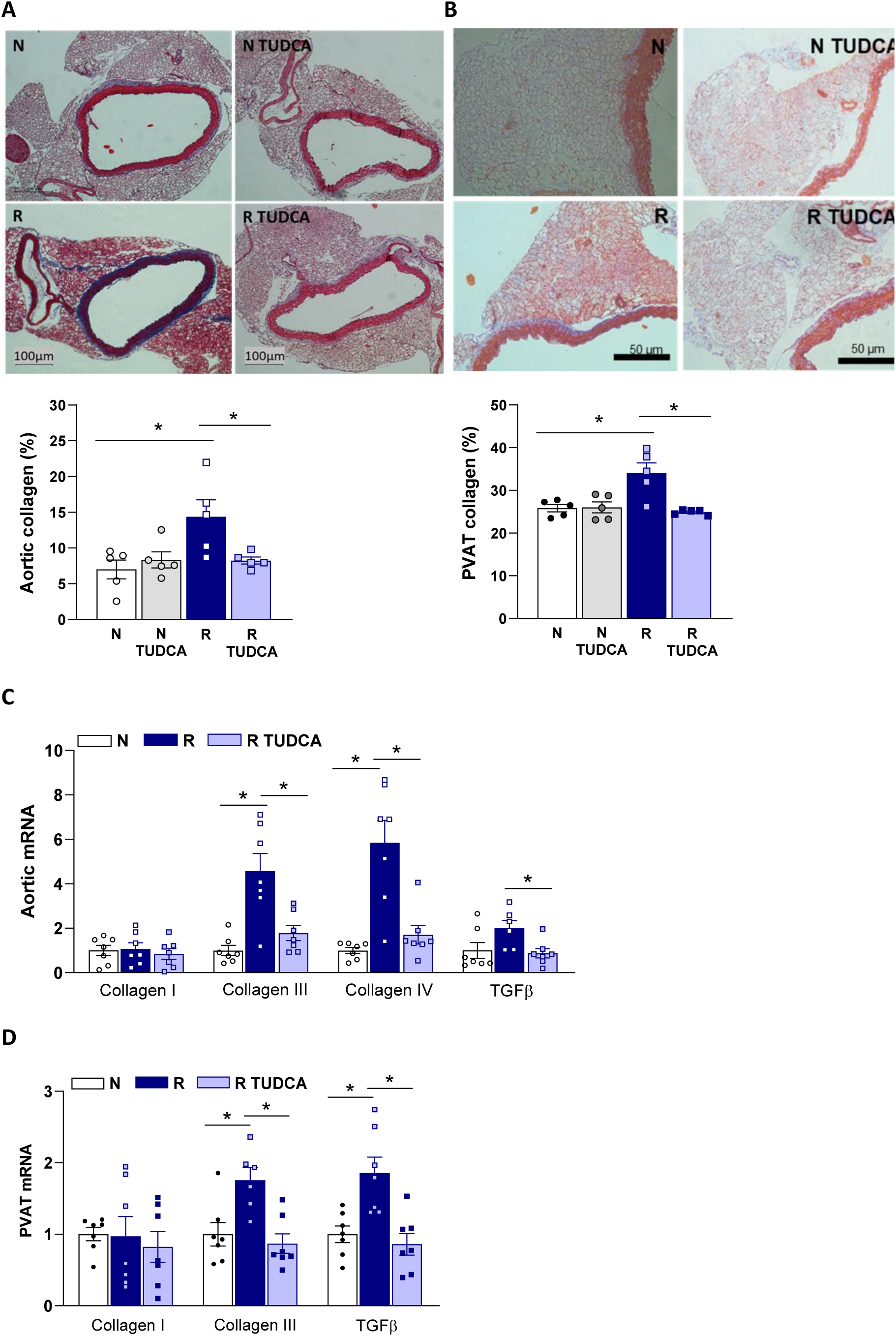
Protein restriction-induced aortic and PVAT fibrosis is prevented by TUDCA treatment. Collagen deposition was assessed by Masson’s trichrome staining and quantified in **(A)** aorta (bar= 100 µm) and **(B)** PVAT (bar= 50 µm) tissue sections from mice on a normoprotein (N) or protein-restricted (R) diet, with or without TUDCA treatment. mRNA expression of collagen subtypes I, III and IV, and TGF-β were evaluated in the **(C)** aorta and **(D)** PVAT. RNA expression was normalized to the internal control gene Cypa (aorta) or Rplp0 (PVAT). Data are presented as mean ± SE. *P<0.05: 1-way ANOVA.

### ER stress inhibition and FXR upregulation following TUDCA treatment in the aorta and PVAT of protein-restricted male mice

Protein restriction upregulated ER stress-related genes in the aorta and PVAT, including GRP78 and activation of the three UPR branches (PERK/ATF4/CHOP, IRE1α/XBP1, and ATF6), which was normalized by TUDCA, although XBP1 was not modulated in the PVAT (Figures 4A-B). TUDCA cell signaling may rely on bile acid receptors including TGR5, S1PR2 and FXR. Here we found reduced gene expression of FXR in both aorta and PVAT of protein-restricted mice, that were upregulated by TUDCA (Figures 4C-D), while TGR5 and S1PR2 gene expression were similar between groups (Figures 4C-D).

**Figure 4:**
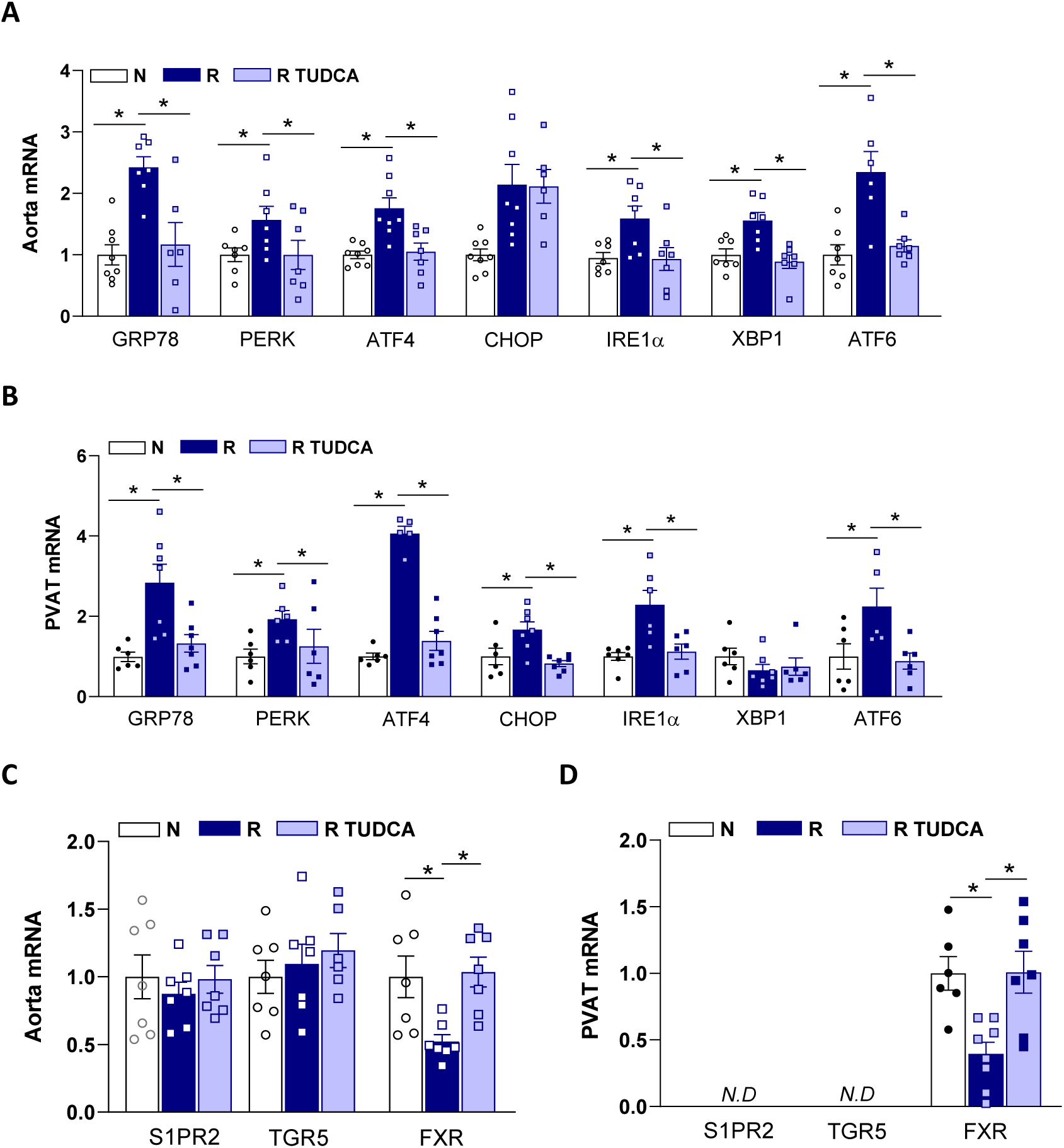
T**U**DCA **normalized endoplasmic reticulum (ER) stress markers and FXR expression in the aorta and PVAT of protein-restricted mice.** mRNA expression levels of ER stress markers GRP78, PERK, ATF4, CHOP, IRE1α, XBP1 and ATF6 in **(A)** aorta and **(B)** PVAT, and mRNA expression of TUDCA receptors S1PR2, TGR5 and FXR in **(C)** aorta and **(D)** PVAT of mice fed a normoprotein (N) or protein-restricted (R) diet, with or without TUDCA treatment. Expression was normalized to the internal control gene Cypa (aorta) or Rplp0 (PVAT) and are presented as mean ± SE. *P<0.05: 1-way ANOVA.

To determine whether the beneficial effects of TUDCA on protein restriction–induced vascular alterations are related to ER stress inhibition, male mice in the R group were treated with another chemical chaperone, PBA. PBA treatment did not alter the reduced body weight or total plasma protein levels in the R group (body weight: R=25±3 vs. R PBA=26±2 g; plasma protein: R=4.5±1 vs. R PBA=5.3±1 g/dL; p>0.05) but significantly reduced the blood pressure (R=124±5 vs. R PBA=98±2 mmHg; p<0.05). PBA also reversed the hypocontractility, impaired relaxation to acetylcholine, and the anticontractile effect of PVAT in the aorta of male R mice (Figure 5A-E).

**Figure 5:**
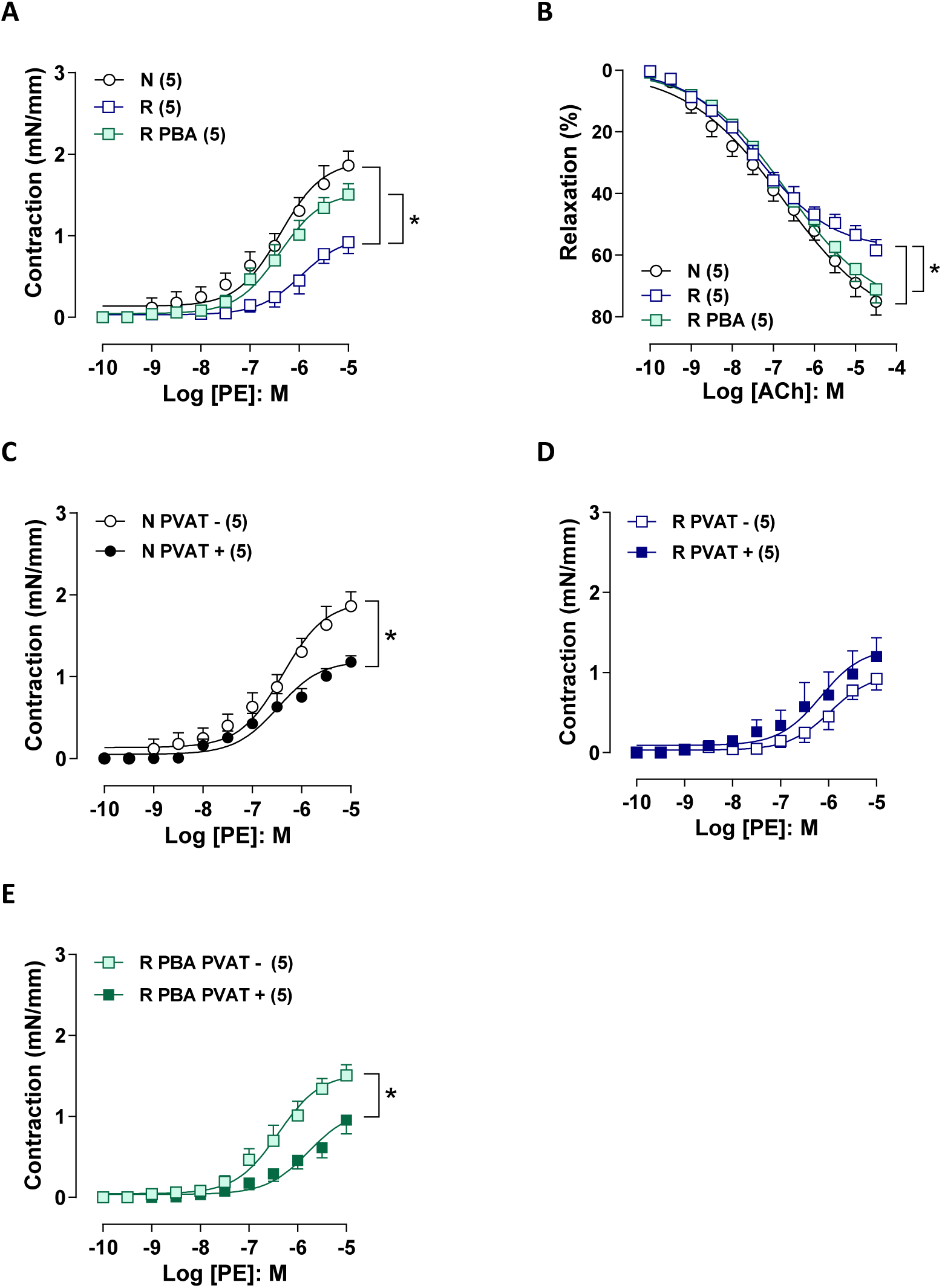
The chemical chaperone 4-phenylbutyrate (PBA) restored aortic contraction, endothelium-dependent relaxation, and the PVAT anticontractile effect in protein-restricted mice. Concentration-response curves for phenylephrine (PE) (A) and acetylcholine (ACh) (B) in aortas without PVAT (PVAT-) in mice fed a normoprotein (N) or protein-restricted (R) diet treated or not with PBA. The effect of PVAT presence (PVAT+) on PE-induced contraction was evaluated (C, D, E). *P<0.05: 2-way ANOVA.

## DISCUSSION

Undernutrition is a form of malnutrition raised since the COVID-19 pandemic in 2020 and with regional differences in prevalence globally. Vascular complications are related to the long-term impact of undernutrition but therapy to reduce the development of cardiovascular disease associated with this condition is poorly explored. TUDCA has emerged as a potential therapy in metabolic disorders (revised in 20). In the present study we found that the treatment with TUDCA restored protein restriction-induced impaired contraction, endothelial and PVAT dysfunction, and vascular and PVAT fibrosis, in addition to reduce blood pressure to baseline levels. Those cardiovascular changes were not accompanied by modification on body weight, total plasma protein and fasting glycemia. These data suggest cardiovascular benefits of TUDCA in the setting of early-life protein restriction.

Our results corroborate previous studies showing that dietary protein restriction during early life predisposes to long-term cardiovascular complications. Increased blood pressure and reduced endothelial-dependent relaxation was demonstrated in arteries of protein-restricted male mice (7; 31). In addition, here we found impaired aortic contractility, PVAT dysfunction and aortic and PVAT fibrosis in response to early-life protein restriction. Interestingly, even protein restriction was modeled in females by reducing body weight, total plasma protein and increasing blood pressure, no changes in vascular reactivity was observed. This suggested a relative protection for protein restriction long-term vascular complications in female sex. This agrees with males being more vulnerable to the adverse effects of postnatal undernutrition including brain and gastrointestinal injury and to coagulopathy (32–34). Ovariectomy impairs vascular contractility, endothelium-dependent relaxation and the anticontractile effect of PVAT (28; 36). But little is known regarding sex hormones and sex chromosomes mediating differences in vascular health impact of early nutrient deprivation.

Small-for-gestational-age children have reduced levels of circulating NO, which correlate with vascular function and higher blood pressure, elevating the risk of vascular disease later in life (37). Early life protein restriction may contribute to this context by reducing eNOS expression and NO production (6). In addition, NO levels are reduced in human endothelial cells exposed *in vitro* to amino acid restriction (31). However, the molecular mechanisms governing vascular alterations in this condition are poorly explored. ER stress has been described as a mechanism of vascular and endothelial injury. Pharmacological activation of ER stress promotes phenotypic switching of vascular smooth muscle cells which may contribute to impaired vascular contractility (38) and impairs endothelium-dependent relaxation in mouse aorta through TGF-β signaling (39). In addition, ER stress inhibition with TUDCA reduced blood pressure and improved endothelium-dependent relaxation and vascular fibrosis in aorta of angiotensin II, DOCA-salt and SHR models of hypertension (39–41). In humans, TUDCA administration prevented endothelial dysfunction caused by hyperglycemia (42) and obesity (43). In line, our findings demonstrated that TUDCA protected from increased ER stress markers, fibrotic factors, collagen deposition, endothelial, and contraction dysfunction indicating relevant vascular benefits for ER stress inhibition in the context of protein restriction.

Treatment with TUDCA increases lipolysis and insulin signaling in brown and white adipose tissue of protein-restricted obese mice (11), suggesting ER stress as a mechanism that may confer higher susceptibility to adipose tissue dysfunction in the context of protein restriction. In our study, TUDCA treatment restored the reduced anticontractile function and lipid content in PVAT of protein-restricted mice, accompanied by normalization of PRDM16 (BAT development marker), PPARγ (adipogenic marker), and leptin/Ob-Rb expression. Reduced PVAT mass is associated with altered vascular reactivity and increased blood pressure (44), and PPARγ deficiency enhances atherosclerosis and macrophage infiltration in the PVAT (45). PVAT ER stress has been suggested as a mechanism of vascular dysfunction as TUDCA was associated with improved vascular function and reduced vascular stiffness in type 2 diabetic mice (46) as well as PBA locally administrated to PVAT reduces atherosclerotic lesion in apoE-/- mice (47). Therefore, normalization of PVAT adipocyte area and derived factors may contribute to improved vascular function in response to ER stress inhibition by TUDCA in in protein-restricted mice. Protein restriction induced a lipodystrophy-like PVAT remodeling, characterized by smaller adipocytes and fibrosis (48). A fibrotic PVAT was recently demonstrated to be associated with aortic aneurism (49) highlighting the therapeutic importance of targeting PVAT remodeling.

The similar effect of TUDCA and PBA support the role of ER stress mediating endothelial and PVAT dysfunction in protein restriction. A limitation of our study is that although we found that TUDCA inhibited the expression of the 3 major UPR cascade it is unclear if a particular cascade contributes to the major vascular effects of protein restriction. Recently, it was demonstrated that IRE1α inhibition mitigate hypertension and vascular remodeling induced by angiotensin II supporting a role for this pathway in the ER stress induced vascular dysfunction (44).

Malnutrition-induced coagulopathy in males was associated with decreased expression of FXR target genes, decreased FXR activation and binding, and decreased production of FXR ligand bile acids (33). Here we found decreased gene expression of this bile acid receptor in aorta and PVAT of protein-restricted mice that was reversed by TUDCA treatment. FXR in adipose tissue enhanced PPARγ-dependent adipocyte differentiation and downregulated proinflammatory cytokines in brown adipose tissue (50; 51). In the vascular tissue, FXR activation decreases vascular inflammation and calcification (52). These findings support a putative protective effect for FXR agonism by TUDCA in the aorta and PVAT. S1PR2 and TGR5 gene expressions were below the limit of detection PVAT and not modified by TUDCA treatment. As FXR agonists have been shown to attenuate ER stress (53; 54), it is possible that this signaling pathway adds benefits to the chaperone activity of TUDCA. The contribution of FXR to the vascular and PVAT effects of TUDCA needs to be addressed in future studies.

In conclusion, vascular alterations induced by long-term protein restriction exhibited sexual dimorphism, with males being more susceptible than females. ER stress was identified as a key mechanism mediating vascular and PVAT dysfunction and fibrosis in the setting of post-weaning protein restriction in males. ER stress inhibition with TUDCA lowered blood pressure and reversed vascular and PVAT remodeling, associated with increased expression of the bile acid receptor FXR. These data suggest TUDCA may represent a promising strategy to mitigate vascular complications associated with long term undernutrition, a neglected world health problem.

### Perspectives

In this study, we demonstrated that treatment with the ER stress inhibitor TUDCA reverses the increased blood pressure, altered vascular reactivity, impaired endothelial and PVAT function, as well as aortic and PVAT fibrosis induced by post-weaning protein restriction. These findings identify ER stress as a mechanistic link between early-life protein restriction and long-term cardiovascular dysfunction, highlighting ER stress modulation as a potential therapeutic target. TUDCA treatment increased FXR expression in both the aorta and PVAT, suggesting that this receptor may act as a modulator of beneficial vascular effects in malnutrition, a possibility that should be addressed in future studies. Our data expand the understanding of the vascular consequences of early-life malnutrition and suggest TUDCA as a potential adjuvant strategy to mitigate cardiovascular risk associated with chronic undernutrition.

## Supplemental Material

Supplemental Methods Tables S1-S3

Figure S1

## NOVELTY AND RELEVANCE

### What Is New?

The ER stress inhibitor TUDCA restored early protein restriction-induced vascular and perivascular adipose tissue (PVAT) dysfunction and remodeling in males.

### What is Relevant?

Male sex appears more susceptible than females to the vascular consequences of early protein restriction. ER stress was identified as a central mediator linking early-life undernutrition to endothelial, vascular, and PVAT dysfunction and fibrosis.

### Clinical/Pathophysiological Implications?

TUDCA may represent a potential adjuvant strategy to mitigate cardiovascular risk associated with early-life malnutrition through ER stress inhibition and FXR signaling.

